# Protein Structural Biology Using Cell-Free Platform from Wheat Germ

**DOI:** 10.1101/375188

**Authors:** Irina V. Novikova, Noopur Sharma, Trevor Moser, Ryan Sontag, Yan Liu, Michael J. Collazo, Duilio Cascio, Tolou Shokuhfar, Hanjo Hellmann, Michael Knoblauch, James E. Evans

**Affiliations:** Environmental Molecular Sciences Laboratory, Pacific Northwest National Laboratory, 3335 Innovation Blvd., Richland, WA, 99354, USA; School of Biological Sciences, Washington State University, Pullman, WA, 99164, USA; Department of Biological Chemistry and Department of Chemistry and Biochemistry, University of California Los Angeles, Howard Hughes Medical Institute, UCLA-DOE Institute for Genomics and Proteomics, Los Angeles, CA, 90095, USA; Department of Bioengineering, University of Illinois at Chicago, Chicago, IL, 60607, USA

**Keywords:** Cell-free protein expression, wheat germ, electron microscopy, cryo-EM, X-ray, structural biology, protein purification

## Abstract

One of the biggest bottlenecks for structural analysis of proteins remains the creation of high yield and high purity samples of the target protein. Cell-free protein synthesis technologies are powerful and customizable platforms for obtaining functional proteins of interest in short timeframes while avoiding potential toxicity issues and permitting high-throughput screening. These methods have benefited many areas of genomic and proteomics research, therapeutics, vaccine development and protein chip constructions. In this work, we demonstrate a versatile and multistage eukaryotic wheat-germ cell-free protein expression pipeline to generate functional proteins of different sizes from multiple host organism and DNA source origins. We also developed a robust purification procedure, which can produce highly-pure (>98%) proteins with no specialized equipment required and minimal time invested. This pipeline successfully produced and analyzed proteins in all three major geometry formats used for structural biology including single particle analysis, and both two-dimensional and three-dimensional protein crystallography. The flexibility of the wheat germ system in combination with the multiscale pipeline described here provides a new workflow for rapid generation of samples for structural characterization that may not be amenable to other recombinant approaches.

## Introduction

Systems Biology seeks to understand genomic and proteomic changes between species or between individual cells to link compositional changes to the genome (or proteome) to the observed phenotype. However, many organisms still have an array of genes and proteins of unknown structure and function while other annotated proteins may have limited homology to proteins in the same class. For example, almost 2500 of the 8000 genes identified in the smallest eukaryote *Ostreococcus tauri* are designated as proteins of unknown function [1, 2]. Understanding the structure, mechanism and function of these unknown proteins is the key to linking information across scales from the individual protein to whole cell/organism. Thus, a better link is needed to connect proteomic output with structural biology pipelines.

Today, a variety of cell-free expression options have emerged as powerful alternatives to laborious *in vivo* methods of protein synthesis to support the growing demand for easy and cost-effective ways for protein production. Depending on the biochemical properties, the origin and the potential use of the proteins to be expressed, several cell-free protein expression systems can be explored and are commercially available today. These are based on *E. coli*, wheat germ, rabbit reticulocyte, *L. tarentolae*, insect and human cell extracts [3–9]. Although not currently commercially available, efficient cell-free lysates can also be prepared from tobacco BY-2 cells, Chinese hamster ovary (CHO) and yeast [10–13]. All cell-free translation systems rely on two major components: 1) a specifically designed vector or a PCR template with your gene of interest, and 2) a cell extract of choice that matches the experimental needs for yield or post-translational modifications. The reagents are supplemented with additional energy sources, amino acids and various cofactor molecules to aid continuous protein synthesis and folding. Key advantages of these systems are the ease of the setup, fast turnaround times, linear scalability for the reaction volumes and amenability to high-throughput screening. Additionally, these platforms allow significant customization as one can directly provide additives to aid solubilization and folding of difficult targets [14–16]. One can also include unnatural amino acids to facilitate labeling and characterization or introduce additional DNA/RNA templates to generate diverse protein hetero-assemblies by co-expression [17–22]. Since there is no need to sustain a living organism, toxic proteins that failed to be produced by cell-based methods can be readily expressed [23]. These numerous benefits are being widely exploited in applications of NMR and X-ray structure determination, functional genomics and proteome research, protein chip construction, therapeutics and vaccine development [17, 23–26].

While thousands of proteins can be manufactured in a high-throughput fashion using the cell-free translation format [26, 27], structural biology primarily requires a supply of specific protein components. Thus, the most critical parameters are high yield and high purity of the protein sample. High–resolution structural studies require at least 50 μg of highly-purified protein for cryo-EM whereas X-ray crystallography can require 1 mg or more. As of today, in comparison with other cell-free systems, *E. coli* cell-free protein production is the most frequently used for the needs of structural biology in X-ray and NMR due to its cost and availability [24]. However, a wheat germ lysate has the highest solubility and the highest translation yields for eukaryotic proteins. For instance, in the high-throughput proteome study, 12,996 human clones (out of 13,364 tested) gave rise to proteins whereas 97.6% of those were detectable in a soluble fraction [26]. In addition, codon optimization is not required for wheat germ, which makes it compatible with various DNA templates from different organisms of both prokaryotic and eukaryotic origins [28]. Despite these benefits, the use of wheat germ cell-free system for structural biology has been primarily limited to NMR studies [17]. There is only one report on the use of this system for X-ray structure determination of PabI protein [29]. However, the purification of PabI from the crude mixture was done based on its heat-resistance properties and this is not applicable for other protein targets.

The Environmental Molecular Sciences Laboratory is a United States Department of Energy user facility that is primarily focused on organisms relevant to bioenergy and the environment such as bacteria, algae, fungi and plants. Using a wheat germ cell-free expression system from CellFree Sciences, invented by the Endo group from Japan [4, 5], we sought to build a cell-free expression pipeline in order to link the systems biology and structural biology capabilities at this user facility while maintaining applicability to a wide-range of organisms and being compatible with all sample geometries used for high-resolution electron microscopy and x-ray structural analysis. Here we describe a multistage pipeline that allows quick screening for protein solubility, quick optimization of expression yield along with purification strategies suitable for high-resolution structural investigations using several target genes from various diverse organisms (bacteria, algae and plants).

## Results

Our workflow included four different scales of protein expression (MINI, MIDI, MAXI and MEGA) to allow screening for solubility and impact of additives (MINI), overall expression levels and purification optimization (MIDI), and scale-up for cryo-EM (MAXI) or X-ray crystallography applications (MEGA) (Figure 1).

**Figure 1.**
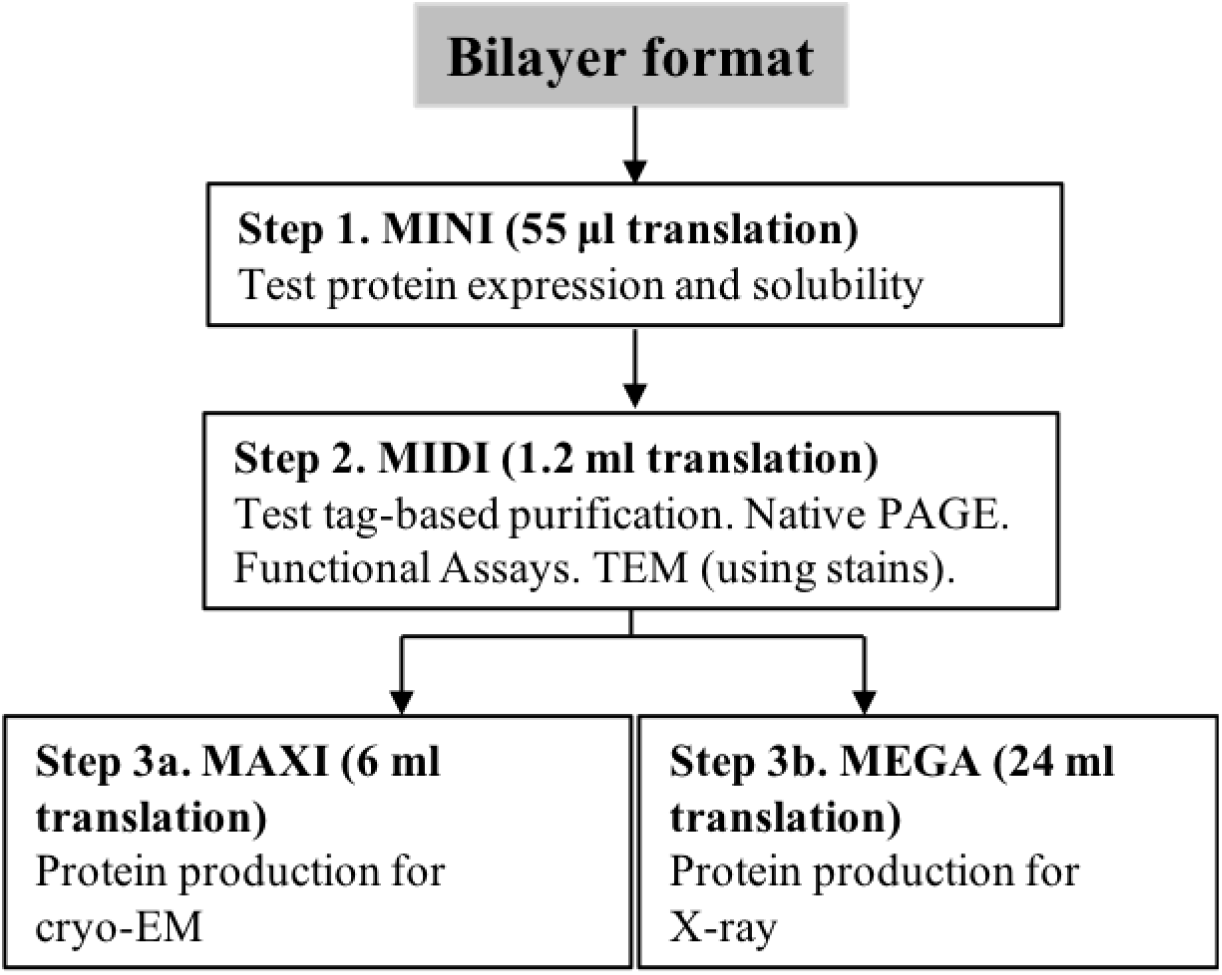
Experimental workflow for the cell-free expression platform used in this study.

### Quick translation trials to determine the protein expression potential (MINI)

PCR templates for cell-free expression can be prepared in short amount of time (compared to construction of vectors) and can therefore be used to quickly test target protein for expression and solubility. Our first stage involves MINI-translation reactions using PCR templates and fluorescently labelled tRNA. These reactions are only 55 microliters in volume allowing for small-scale testing of various conditions with minimal expenditure of reagents. The PCR template requires just three elements: a SP6 transcriptional promoter, an E01 translational enhancer sequence and a 3’-UTR for better protection of mRNA against degradation (Figure 2A) [30]. To incorporate those elements, a two-step PCR using a split primer design is suggested [30, 31]. In the first round of PCR (PCR1), the gene of interest is amplified using a pair of gene-specific primers, where a reverse primer is designed to bind at ~1.4-1.6 kB away from the termination codon. At the second round of PCR (PCR2), the SP6 promoter and the E01 sequence are introduced through two forward SPU and deSP6E01 primers. The final PCR construct serves as the template for transcription and MINI-scale translation. To create a universal approach in handling DNA from different sources, we carried out all PCR reactions using the Q5 DNA polymerase from NEB, a high-fidelity enzyme with ultra-low error rates and suitable for low, medium and high GC contents of DNA template [32]. In addition, the Q5 enzyme demonstrates better capacity to produce long amplicons.

**Figure 2.**
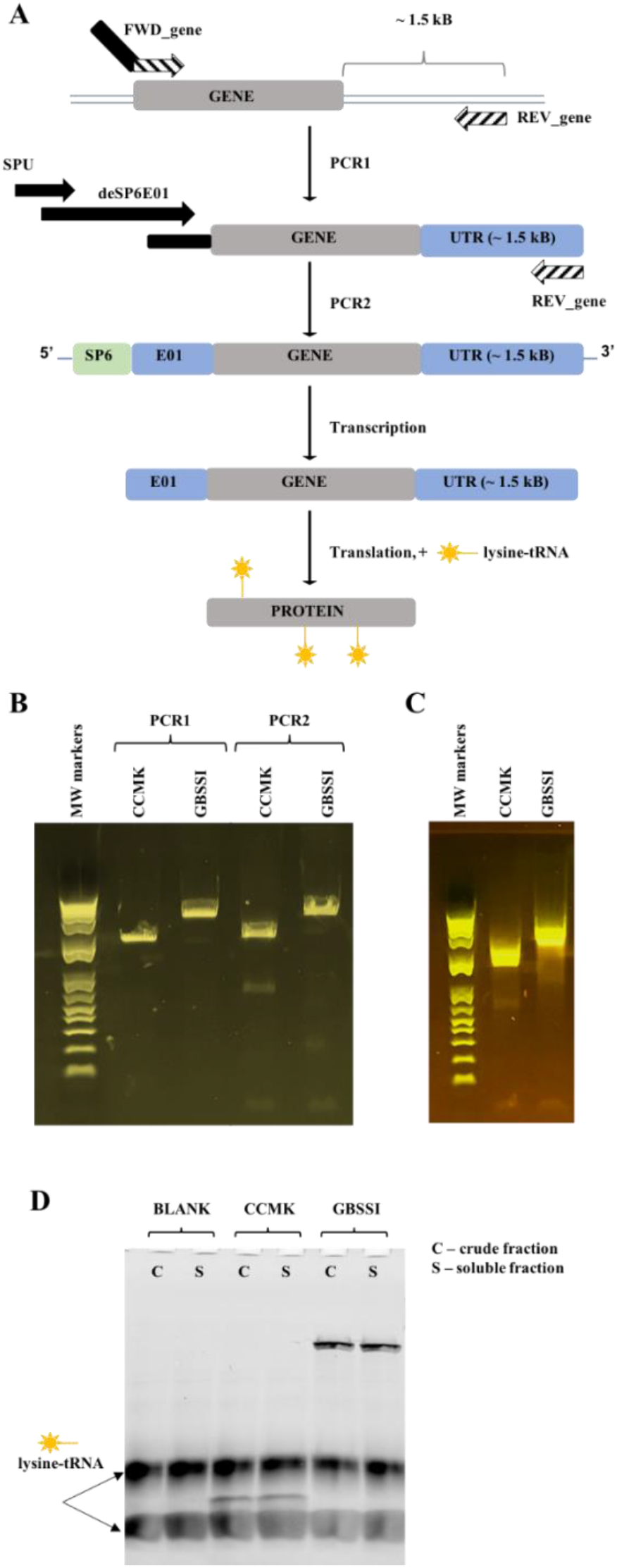
Expression trials using PCR templates. A. Schematic of two-step PCR template preparation for CellFree Sciences wheat germ expression kit followed by transcription and translation in the presence of the FluoroTect Green_Lys_ tRNA. Elements shown are the SP6 promoter (SP6), the translational enhancer sequence (E01) and the untranslated region (UTR). The **SPU** and **deSP6E01** are the names of the primers used. B. The detection of PCR products on agarose. C. The detection of mRNAs on agarose. D. SDS-PAGE gel showing the expression of fluorophore-containing translation products. BLANK is a negative control translation which has no PCR template added to the reaction mixture.

PCR-based expression trials were conducted on two genes of diverse origin and size: carbon dioxide-concentrating mechanism protein (CCMK, 10.6 kDa) from *Prochlorococcus marinus* and granule found starch synthetase (GBSSI, 62 kDa) from *Ostreococcus tauri*. The PCR template for CCMK was successfully amplified from a commercially purchased cDNA clone while the GBSSI gene was specifically amplified directly from the genome of *O. tauri* (Figure 2B). The subsequent PCR2 reactions yielded the desired amplicons and were good templates for high-yield production of full-length mRNAs, which were further introduced into MINI-translation mix (Figure 2C, 2D). For quick detection of the protein product, all MINI-translation reactions were supplemented with the fluorophore-labelled lysine-charged tRNA (FluoroTect^™^ Green_Lys_) for incorporation of fluorophore-tagged lysine into the synthetized protein. The use of the fluorophore-tagged lysine allows detection of picomolar levels of protein with SDS-PAGE, which aids in screening. Both CCMK and GBSSI proteins were expressed efficiently using this cell-free format and displayed high solubility. Although not seen in the current experiment, it is important to note that this quick screening procedure can also useful in detection of premature termination in the crude extract, which may indicate needs for additional supplementation or codon optimization.

### Nested PCR to fish out the “hard-to-amplify” genes from genomic DNA

As described above, the GBBSI protein was easily amplified specifically from the genome of *O. tauri*. However, direct amplification of genes of interest from genomic DNA can often fail for many reasons. The complexity and the size of the genome, repetitive regions, non-optimal primer design are among some of the defining factors [34]. We encountered such failures for the glutamine synthetase (GS, 75 kDa) and pyruvate, phosphate dikinase (PPDK, 100 kDa) genes that we tried to amplify directly from the genome of *O. tauri*. To increase our chances for success, we employed a nested PCR strategy [35]. The idea of nested PCR (Figure 3A) is that one set of primers is used to first amplify a larger portion of the genome containing your gene of interest. Depending on the genome complexity and size, it is common that the desired band will not even be visible due to nonspecific priming events. Nevertheless, the resulting PCR library exhibits much less complexity and is further subjected to a second round of PCR using a pair of the gene-specific primers to amplify the target gene alone. The latter approach significantly increases the specificity of DNA amplification, and thus it has been used extensively for viral detection in clinical samples and in mutation screening assays [35–37].

**Figure 3.**
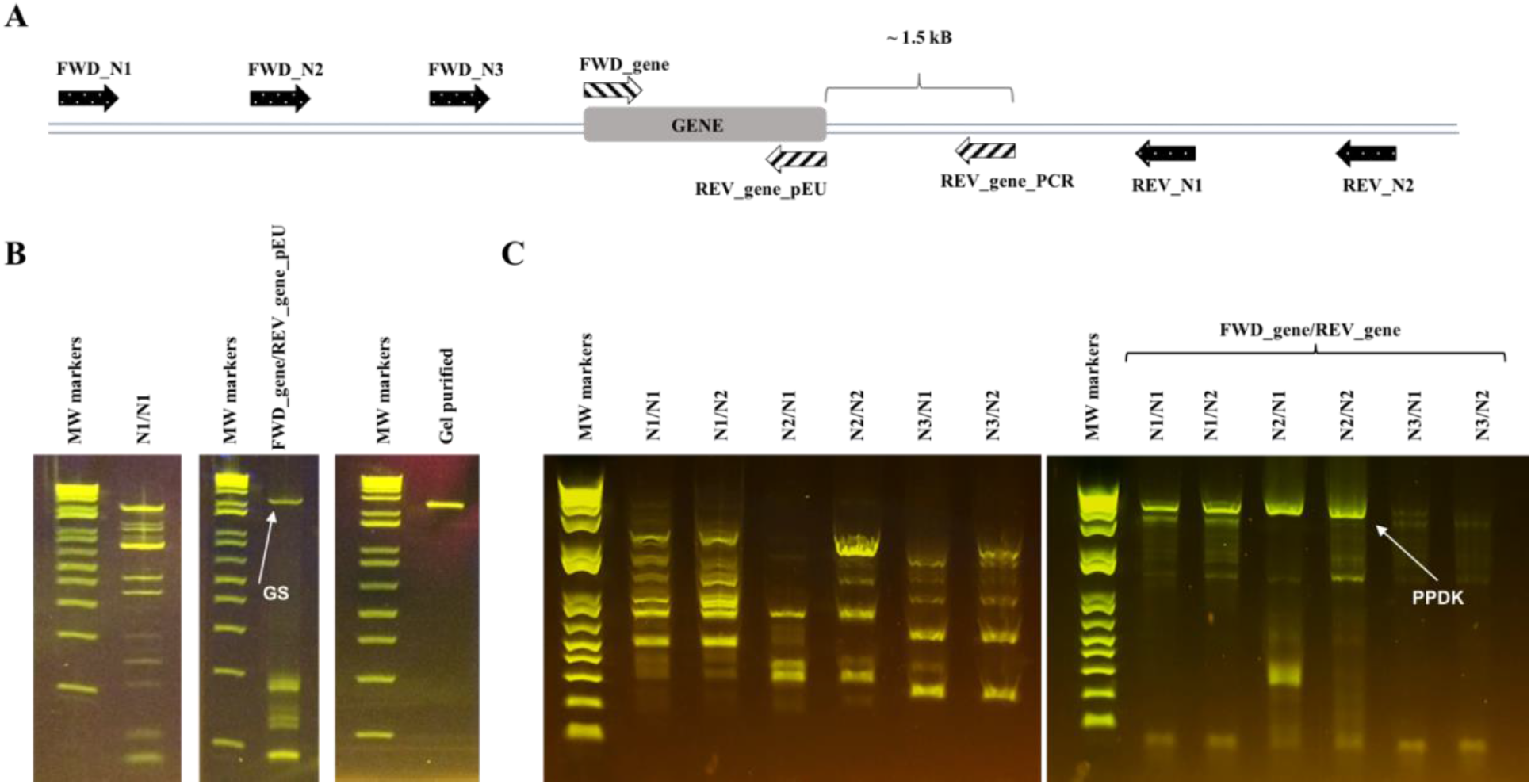
Nested PCR approach for “hard-to-amplify” genes from the genomic DNA. A. A general primer design for nested PCR. First round of PCR uses primers denoted with _N and filled with black. They should bind 0.5kB-1.5kB upstream and downstream from the desired gene. The second round of PCR employs two on-gene primers: FWD_gene and REV_gene_for_pEU primers for insertion in pEU plasmid or FWD_gene and REV_gene_PCR for PCR template. B. Nested PCR amplification of Glutamine Synthetase (GS) gene from the genome of *O. tauri*. C. Nested PCR amplification of Pyruvate, Phosphate Dikinase (PPDK) gene from the genome of *O. tauri*.

Use of only one pair of nested FWD_N1 and REV_N1 primers proved sufficient to amplify GS (Figure 3B). A wide range of PCR products is observed in the first round, while at the second round only one higher molecular weight DNA band of the desired length emerged. The DNA band was further excised, gel-purified and used in the Gibson Assembly with the linearized pEU plasmid to test for vector-based cell-free translation discussed in the following sections. The specific amplification was confirmed by the sequencing of clones. We also tested this approach to amplify another “difficult” gene - PPDK. The GC content reaches 73% in the upstream region and N-terminal portion of the PPDK gene, which gives poor prognosis for its amplification. Indeed, multiple efforts to amplify PPDK directly failed. Therefore, in the nested primer design, we also included an additional nested FWD_N3 primer in order to generate two additional PCR libraries (Figure 3C). Quite diverse PCR populations were generated in the first round of PPDK amplification. They were further subjected to a second round of PCR using gene-specific primers, and four out of six libraries generated similar higher molecular weight products of expected to 4.4 kB size. Subsequent purification and sequencing of these products confirmed that PPDK gene was successfully amplified from N2/N2 library.

### pEU vectors, protein synthesis and purification (MIDI)

If PCR screening trials were successful, the gene of interest was further cloned into our modified pEU vector. For high-yield applications, the optimal expression template is the pEU vector [33], which was modified in house to include a 3XFLAG tag resulting in vectors (pEU-3XFLAG-gene, pEU_3XFLAG_gene_10His, pEU_gene_3XFLAG). Every newly-produced plasmid was also tested at a MINI-scale to access the quality of produced mRNA and protein. The quality of the plasmid itself was found to be critical for obtaining high yields of mRNA and thus high yields of protein sequentially. An example of low quality mRNA can be found in the Supplemental Material (Figure S1). To avoid the latter, the plasmid should be free of RNases, which can originate from plasmid preparation kit itself. However, we usually find that the main cause of bad quality RNA is the plasmid mechanical shearing such as nicking during general plasmid purification procedures. This nicking is not critical for many other applications, but, for *in vitro* transcription reactions, premature runs-off are observed. Therefore, during plasmid productions, gentle cell resuspension and minimal vortexing of DNA are warranted.

The main advantage of cell-free synthesis is the linear scalability of the reaction volumes where reactions can be run as MINI-expression trials to determine gene suitability or scaled-up for high-resolution structural studies (Figure 1). If the plasmid passed the MINI-scale stage, we moved to a MIDI-scale production. The goal here is to determine if the tag attached to either the 3’ or 5’ end of the protein is fully available for efficient binding. As mentioned above, CCMK and GBBSI were found to be good candidates for the cell-free production using PCR screening. Therefore, both genes were cloned into pEU plasmids and tested for protein synthesis and purification using two different tags: 6xHis and 3XFLAG. Initially, the CCMK protein was fused with 6xHis tag and purified using HIS-SELECT magnetic beads (Figure 4A). Naturally, the wheat-germ extract has many endogenous proteins which will nonspecifically bind to nickel, therefore the wheat germ extract optimized for His-tag purification was also employed. A single band of CCMK of desirable purity is clearly seen in the final fraction, which was later used for 2D crystallization trial (discussed later).

**Figure 4.**
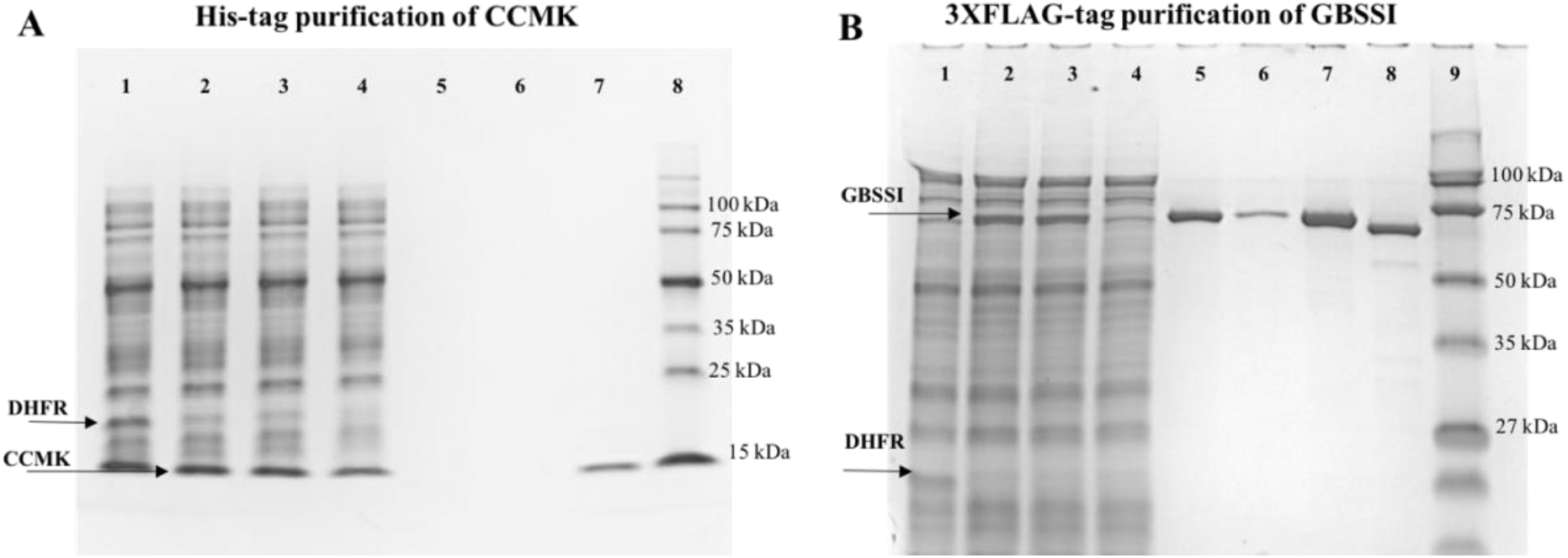
Protein synthesis and purification. A. SDS-PAGE gel, showing the expression and purification of CCMK protein using 6xHis-tag. Lane 1, control DHFR expression; Lane 2, crude mixture; Lane 3, soluble fraction; Lane 4, flow-through; Lane 5, elution fraction 1; Lane 6, elution fraction 2; Lane 7, elution fractions 1 and 2 combined and concentrated; Lane 8, MW markers. B. SDS-PAGE gel of expression and purification of GBSSI using 3XFLAG-tag. Lane 1, control DHFR expression; Lane 2, crude mixture; Lane 3, soluble fraction; Lane 4, flow-through; Lane 5, elution fraction 1; Lane 6, elution fraction 2; Lane 7, elution fractions 1 and 2 combined and concentrated; Lane 8, 3XFLAG tag removed; Lane 9, MW markers.

Despite the use of His-tag optimized wheat germ extract, the desired purity of CCMK was achieved at the expense of the product yield where 50 mM imidazole washes were employed to remove nonspecifically bound proteins. The same purification procedure with 10 mM imidazole washes generated much more material. However, a significant fraction of endogenous proteins was recovered as well unsuitable for our applications (Figure S1B, Lane 7). Ideally, one can try a range of imidazole concentrations in order to find the optimal value. However, to avoid optimization trials in this regard, we decided to employ a 3XFLAG-based purification scheme. The GBSSI protein was first fused with 3XFLAG sequence on its N-terminus and was tested in the purification trial using ANTI-M2 monoclonal magnetic beads. Expressed GBSSI showed good binding to the beads, as indicated by the full disappearance of the expressed protein band in the flow-through fraction (Figure 4B, Compare Lanes 3 and 4). Beads were washed, and then two competitive elution reactions with 3XFLAG peptide were followed to release the protein from the matrix. Both elution fractions had high amounts of released protein (Figure 4B, Lanes 5 and 6). Purified protein appeared to be full-size, homogenous and free of other contaminating proteins coming from the extract itself (Figure 4B, Lane 7). Using enterokinase, the 3XFLAG tag was then removed with >93% efficiency (Figure 4B, Lane 8). The same purification procedure has been tested on many other proteins which some include GS, PPDK, Sieve Element Occlusion protein from *M. truncatula* (SEO1, 75 kDa) [38, 39] and Pyridoxal 5’-Phosphate Synthase-Like subunit (PDX1.2, 34 kDa) from *A. thalianaI* [40]. All SDS-PAGE gels for these proteins are included in the Supplemental Material (Figure S2A). High purity of each final sample was achieved along with high protein recovery. Only PDX1.2 has a somewhat reduced binding to the beads due to its tight structural fold (demonstrated in the next section). It is worth noting that the major advantage of using the 3XFLAG tag is that we observed a complete lack of nonspecific binding. As the result, no optimization is required, and a general one-step purification “bind-wash-elute” protocol can be used for all samples.

### Diverse structural biology applications of cell-free produced proteins

Once we had an isolated protein in hand (tagged or untagged), next step was to verify its activity, assembly state and identify if it is a suitable candidate for high-resolution structural studies. At MIDI-scale, the general yields were determined to be around 20 to 30 μg at 0.2-0.3 mg/ml concentrations. These protein amounts are sufficient for various functional assays, Native PAGE electrophoresis, and preliminary TEM imaging characterization. The structure of CCMK from *P. marinus* has been already available through X-ray crystallography (PDB ID: 4OX8), and the native protein was known to pack functionally as hexamers. Crystallization conditions using a nickelated lipid monolayer were previously determined for a CCMK homolog from *Synechocystis* PCC6803 [41]. Therefore, cell-free expressed and purified CCMK (6xHis tagged) in this study was subjected to a similar 2D crystallization trial. From the first attempt, initial 2D crystal patches were formed and analyzed by TEM imaging (Figure 5A). Negatively-stained CCMK proteins were clearly visualized as hexameric structures, mimicking the packing in the previous study [41]. Image processing of the collected EM micrographs revealed a lattice with reflections extending to 13Å, limited by the negative stain itself. This 2D crystal lattice was able to be directly overlaid by the known atomic structure of *P. marinus* CCMK (RCSB: 4OX8) [42] verifying that the expected structural fold has been adopted by the cell-free synthetized CCMK protein (Figure 5B).

**Figure 5.**
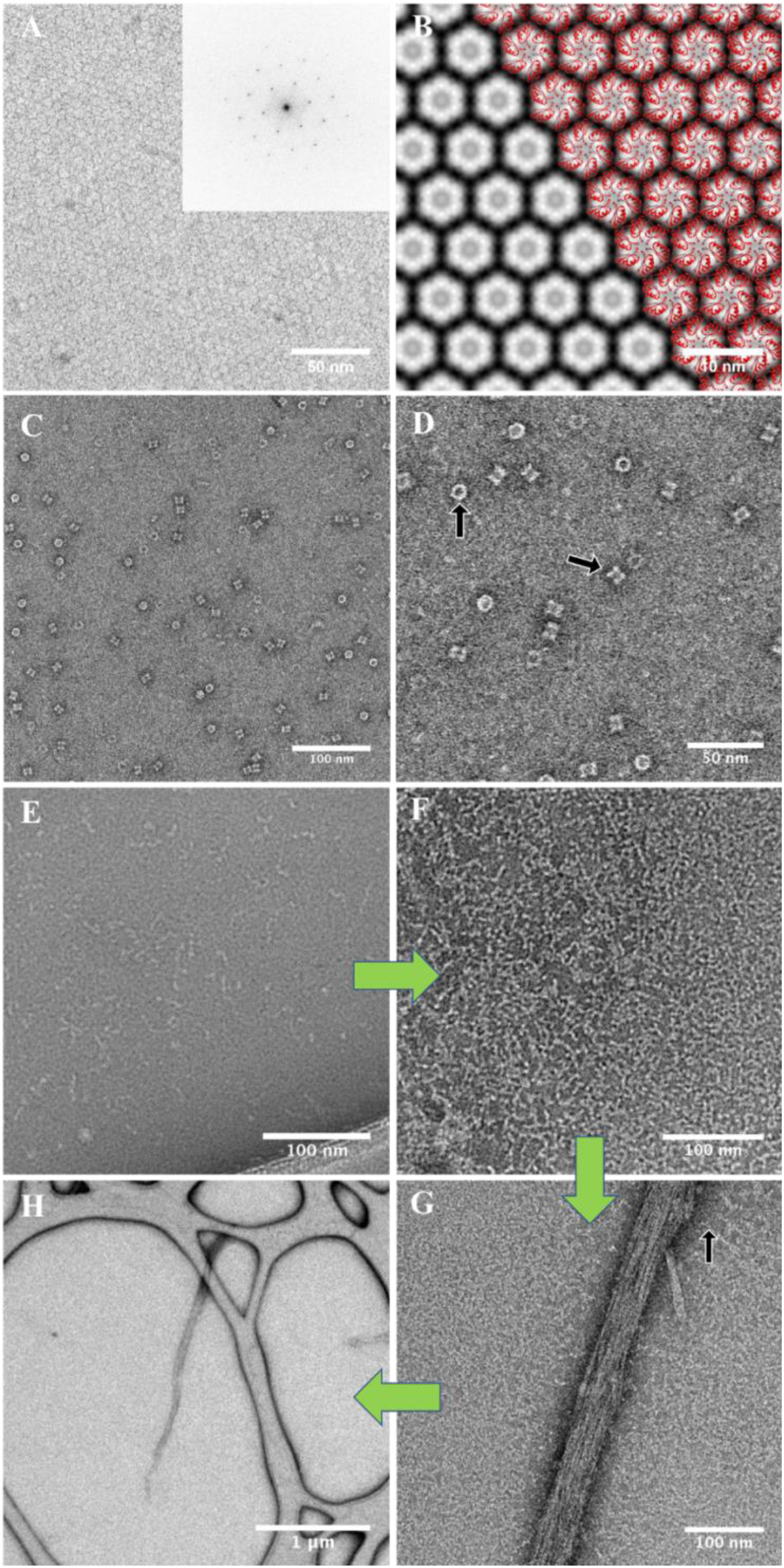
The applications of cell-free produced proteins in electron microscopy. A. STEM image of negatively-stained 2D crystals of CCMK protein (inverted contrast). B. The image from (A) has been processed using a temperature factor of −100, and an IQ cutoff of 2 for the final map. The crystal exhibits P6 symmetry and a unit cell of a=b=57Å and γ = 120°. C,D TEM images of negatively stained PDX1.2 protein complexes. Top- and side-view orientations of the complex are depicted by black arrows. E-H TEM images of negatively stained SEO1 proteins. The individual fibril unit is indicated by a black arrow.

The rest of purified proteins (GBSSI, GS, PPDK, PDX1.2, SEO1) have been initially tested by Native PAGE electrophoresis to gain insight about their potential structural organization and assembly state (Figure S2B). GS and PDX1.2 proteins were found to form defined higher-molecular weight complexes with the sizes of 500 to 600 kDa. TEM images, collected for PDX1.2, showed a two-ring stacked architecture of the complex (Figure 5C,D). No aggregation events or partially disordered complexes were seen, which suggests that the population is quite homogenous where the average diameter of formed complexes was 12 nm. The protein structure closely resembles the structural arrangements of similar PDX1.1 and 1.3 proteins, which form two-ring dodecamers [43]. As the result, PDX1.2 protein seems to form proper structural arrangements and would be a good candidate system for single particle analysis studies by cryo electron microscopy (cryo-EM) after scale-up expression.

We also explored expression of SEO proteins from *Medicago truncatula*, which are known to form large protein bodies, called forisomes [44]. These spindle shaped bodies achieve tens of micrometers in length and look like elongated needles, which swell and deform upon Ca^2+^ introduction. This swelling/deformation is reversible, and it is a critical mechanism in sieve tube flow control [45]. Natural forisomes consist of several members of SEO protein family [38, 39]. However, SEO proteins were never analyzed individually at the structural level. As seen in Native PAGE, cell-free produced SEO1, with molecular weight of 75 kDa, runs higher than 480 kDa control protein complex (Figure S2B). TEM images of SEO1 proteins show that they exist as individual fibrils at initial stages (Figure 5E-H). Upon 3XFLAG removal, SEO1 proteins form even higher order molecular weight structures (Figure S2B). Additional incubation of SEO1 (in Ca^2+−^ free environment, 10 mM EGTA) results in the formation of micrometer long strands, clearly formed through individual fibril assembly (Figure 5E-H). This is consistent with their functional roles *in vivo*, where they participate in the molecular assembly of large bodies. This is the first demonstration that such protein bodies can be generated *in vitro*.

### Scaling-up for cryo-EM (MAXI) and X-ray (MEGA) structural approaches

Due to significant instrumental advances in the electron microscopy such as the development of direct electron cameras and phase plates [46, 47], cryo-electron microscopy (cryo-EM) holds a promise to be major imaging technology to study the structures of biological molecules in its native environment. However, the success of the cryo-EM experiment is also defined by the quality of protein sample such as its purity and homogeneity and typically requires about 100 micrograms of sample for screening and data collection.

Depending on the outcome of the above MIDI-scale tests, the protein synthesis can be further scaled up for high-resolution cryo-EM and X-ray crystallography characterization. Since the negative stain imaging of PDX1.2 showed a fairly homogenous sample, a MAXI scale cell-free expression was performed to generate suitable protein for cryo-EM. While the current 2D class averages and initial 3D volume are limited in resolution due to the conventional microscope used and available for this work (Figure 6A,B), experiments utilizing new state-of-the-art cryo-EM instrumentation are ongoing and will be the focus of a separate paper. Nevertheless, this work demonstrates the ability of this pipeline to generate ample yields for cryo-EM studies especially since the PDX1.2 protein used for cryo-EM validation had the lowest yields of any of the proteins tested.

**Figure 6.**
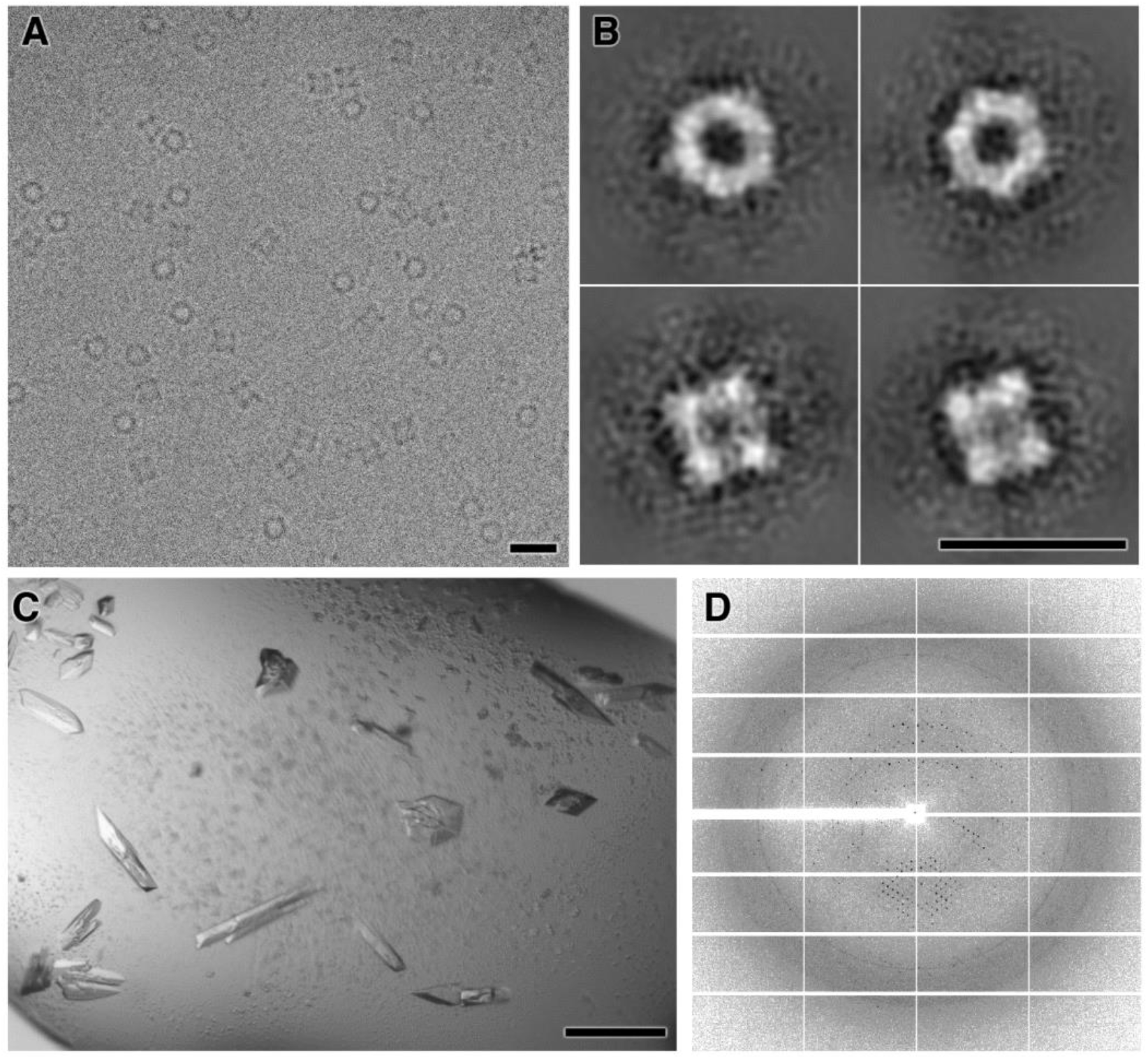
Cryo-EM and X-ray applications of cell-free synthetized proteins PDX1.2 and GS. A. Raw cryo-EM image of PDX1.2 particles distributed in vitreous ice and adopting random orientations. The scale bar corresponds to 20 nm. B. Single particle class averages of cryo-EM dataset from (A) showing class averages for the top, side and 2 intermediate views. A double stacked hexameric ring geometry is evident. The scale bar corresponds to 20 nm. C. Optical image of 3D microcrystals for GS. The scale bar corresponds to 50 um. D. X-ray diffraction pattern collected from the crystals seen in (C).

Finally, we sought to test if the same pipeline could generate enough protein material of good quality for 3D crystallization screening. As other GS classes from cell-based recombinant expression have been successfully crystallized previously [48, 49], we therefore performed a MEGA-scale synthesis of the GS complex and generated enough material (~10 mg/ml, 100 μl) for 3D crystallization trial screening. Several conditions were found suitable for the crystal growth and, without any additional optimization, diffractions up to 5.5 Å were obtained (Figure 6C,D).

## Conclusion

The preparation of a wheat germ cell lysate is a tedious and a quite complex procedure [4], therefore the use of established commercial products is often recommended for obtaining reproducible results [28]. As of 2018, wheat germ cell-free expression (WGCF) kits are available from two manufacturers: CellFree Sciences and Promega. To minimize sample handling, the Promega WGCF kit offers a coupled setup where transcription and translation are run simultaneously. The CellFree Sciences WGCF kits have transcription and translation processes rather decoupled where each step is optimized individually to maximize the yield. If one needs to use an additive for the translation or change the reaction temperature, the latter setup offers a benefit of not interfering with the transcription stage. Due to the above-described reasons and a wider selection of the reagents for a MINI/MIDI or MAXI/MEGA-protein synthesis, we opted to use the commercial system from CellFree sciences.

While the use of the specialized pEU vector is preferred to obtain higher yields of the given protein in this cell-free format, the most time-consuming steps are still the cloning of the respective gene, purification of the plasmid product, its subsequent sequencing and scale-up preparation. While this is the final route to get high quantities of the desired protein product, it is always beneficial to know in advance if the protein is a good candidate to be expressed in this system and is worth the time invested. Fortunately, cell-free expression with somewhat reduced protein yields can be performed using PCR templates. The latter offers an ideal rapid screening to explore the protein potential to be expressed in a soluble and active form. Based on our experiences in expressing a variety of proteins for different applications, we had a very high probability of producing good quality proteins from different organisms using this system.

In combination with the wheat germ cell-free extract, this work presents a useful methodology for a multiscale pipeline covering all stages from screening potential candidates of protein synthesis to final production of micrograms quantities of a highly pure and tag-free protein. We demonstrate how to screen potential protein candidates using two-step PCR, which is suitable for the “easy-to-amplify” genes either sourced from the host vectors or directly from genomic DNA. For the “hard-to-amplify” genes, we devised a nested PCR strategy. Then we demonstrated a vector-based expression and optimized purification pipeline to generate a tag-fused or tag-free protein.

We show that the developed 3XFLAG purification protocol allowed high protein recoveries with purities reaching 98%, and it was found to be our preferred method for preparation of all protein samples. The only exception was to use a His-tag based purification for the 2D protein crystallography method which relies on a Ni-NTA functionalized lipid monolayer. Nevertheless, His-tag purification can be considered a good alternative strategy. In terms of yield, 0.6 mg per ml wheat germ extract has been generally obtained in the bilayer format used here. Not demonstrated here, but for large scale productions, impressive protein yields up to 20 mg per ml WG extract can be achieved by a two-chamber dialysis or a “filter-feed” method using dedicated robotic synthesizers [28]. While we show that such robotic platforms are not necessary, if available at a nearby institution, they could be used to replace the MEGA scale expression using lower reaction volumes at potentially lower cost. In terms of structural function/fold, cell-free produced proteins in this study exhibited expected structure and assembly state in comparison with similar proteins produced and characterized by other means.

The generated protein products are compatible with three major protein sample formats for high-resolution structural studies: (1) single particle analysis, (2) 2D crystallography and (3) 3D macromolecular crystallography. Obviously, the developed pipeline is applicable to any science area where interests lie in expressing of proteins in high yields and purities without the use of any specialized equipment and minimal time invested. Due to the fact that the cell-free expression platforms are gaining significant popularity, we believe that these comprehensive experiments will be useful for a wide range of scientific audience interested in exploring these systems especially as this platform can be replicated without major investments in instrumentation.

## Materials and Methods

### DNA template construction

PCR reactions were carried out using the Q5 Hot Start High-Fidelity 2X Master Mix from New England Biolabs. NEB Tm Calculator (http://tmcalculator.neb.com/#!/) has been used to assess annealing temperatures. DNA primers were acquired from either IDTNA or Invitrogen. The pEU vector (pEU-MCS-TEV-HIS-C1) was from CellFree Sciences. Gene sequences were either amplified from genomic DNA from *Ostreococcus tauri* or *Arabidopsis thaliana (gifts from* M. Knoblauch and H. Hellmann labs), or purchased as synthetic genes from GENSCRIPT. To obtain genomic DNA from *O. taur*i, the DNeasy Plant kit from QIAGEN has been used for extraction but tissue homogenization procedures were omitted. Genomic DNA was further ethanol precipitated and re-suspended in water. The quality of genomic DNA and PCR products was assessed using the E-gel Electrophoresis System (E-gel pre-cast 1% EX Agarose gels with SYBR safe along with E-gel 1kB Plus DNA Ladder) from Invitrogen. Gel purification of DNA was carried out with the help of Zymoclean Gel DNA recovery kit from Zymo Research as instructed. Zymoclean-purified DNA was additionally desalted using the Bio-Rad mini spin columns, pre-filled with the Bio-Gel P-6DG soaked in water. For cloning in the pEU vector, the Gibson Assembly Master Mix from New England Biolabs was used. The linearized pEU vector and the insertion fragment containing the desired gene were prepared by PCR. These fragments were further gel-purified and quantified with NanoDrop 2000c from ThermoFisher. As an example of Gibson reaction, linear pEU plasmid (2 μl, 20 ng/μl) was mixed with linear gene PCR template (3.8 μl, 15 ng/μl), water (4.2 μl) and 2X Gibson Assembly Mix (10 μl) and incubated at 50°C for 1 hour in the thermocycler. The assembled product (2 μl) has been used to transform an NEB10 *E. Coli* supplied with the Gibson Assembly kit. Transformed *E. Coli* was later plated on a carbenicillin-containing agar plate and grown at 37°C overnight. Several individual colonies were then selected and grown overnight in several milliliters of LB broth supplemented with 100 mg/ml carbenicillin. The QIAGEN Plasmid Mini Kit was then employed for plasmid extractions, which were then sent for sequencing to Eurofins Genomics or MCLAB. Successful clones were later grown in larger quantities of LB broth (with carbenicillin) and harvested using the QIAGEN Plasmid Maxi kit. To note, a 3XFLAG tag was initially introduced through PCR primers in the first construct to generate pEU_3XFLAG_LA_gene. For all subsequent cloning experiments for other genes, the obtained pEU_3XFLAG_LA_gene vector was used as a template for other constructs. The general sequence design of pEU vectors can be found in the Supplemental Material (Figure S3).

### Cell-free expression

Protein synthesis was carried out using Wheat Germ Protein Research Kits (WEPRO7240 and WEPRO7240H) from CellFree Sciences as suggested by the manufacturer. All translation reactions were carried out in the bilayer format where total translation volume accounts for the SUB-AMIX layer. For MIDI-scale plasmid-based protein expression (1.2 ml total translation volume), 5 μl of pEU plasmid (1 μg/μl) was mixed with 10 μl of 5xTranscription Buffer, 5 μl of 25 mM NTP mix, 0.625 μl of RNase inhibitor and 0.625 μl of SP6 polymerase and incubated at 37°C for 3-4 hours with gentle shaking. Upon completion, 1 μl of the transcription mixture was loaded on the E-gel precast agarose gel (Invitrogen) to assess the mRNA integrity. Then 50 μl of transcription reaction were mixed with 50 μl of wheat germ extract (WEPRO7240) and 4.25 μl of 1 mg/ml creatine kinase by a pipette, and the entire mixture was then transferred to the bottom of the individual well of a standard 24-well plate, prefilled with 1.1 ml of translation buffer (1x SUB-AMIX SGC). To prevent evaporation, wells were sealed with Parafilm. These reactions were carried out for at least 20 hours at 15 °C with no mixing at Thermomixer C from Eppendorf. At the end, the contents were gently mixed by a pipette and centrifuged at 20,000 g for 15 min. The supernatant fraction was then buffer-exchanged to TBS (50 mM Tris-HCl, pH 7.5, 150 mM NaCl) using pre-equilibrated in TBS Zeba Spin Desalting Columns (5ml) from Thermofisher (this step is critical to remove DTT) and further subjected to the protein purification procedure of choice. For MAXI- and MEGA-scale plasmid-based protein expression (6- and 24 ml translation volumes), all reagents were scaled linearly, and a 6-well plate was used instead.

For PCR-based protein expression (MINI-scale, 55 μl translation), the following changes have been implemented in the transcription and translation protocols: 1) for transcription, 0.5 μl of the PCR product (2 μg/μl) was mixed with 4.5 μl of the Transcription premix and left incubated at 37C for 4 hours, 2) for translation, 2 μl of the resulting mRNA was mixed with 0.5 μl of FluoroTect GreenLys and 2.5 ul of the wheat germ extract (WEPRO9240) and transferred under 50 μl of the translation buffer (1x SUB-AMIX SGC), placed in the 96-well half-area plate. The FluoroTect Green_Lys_ in vitro Labeling System was purchased from Promega. The translation was carried out overnight and the protein products were analyzed by SDS-PAGE.

### Protein purification using a His-tag

Purification using a His-tag has been carried out using HIS-Select Nickel Magnetic Agarose beads from Sigma-Aldrich in combination with the MagneSphere technology Magnetic Separation Stand from Promega. Per 1.2 ml translation (MIDI-scale), 200 μl of gel suspension (100 μl of packed gel) was used. The needed volume of gel suspension was first equilibrated in 50 mM Tris, pH 8.0, 300 mM NaCl, 50 mM imidazole buffer and then combined with the DTT-free translation mixture. The binding was carried out for 1 hour at room temperature with shaking on the ThermoMixer C (~1000 rpm). Using the magnetic stand, the supernatant was removed and discarded, and the beads were then washed 3 times with 1 ml of 50 mM Tris, pH 8.0, 300 mM NaCl, 50 mM imidazole wash buffer. To elute the bound protein, 0.5 ml of 50 mM Tris, pH 8.0, 300 mM NaCl, 500 mM imidazole elution buffer was applied to the beads, and the mixture was left shaking for 30 min. Then, the supernatant fraction was collected, and the elution procedure was repeated one more time. Then the elution fractions were combined, concentrated using 3kDa Amicons from EMD Millipore and buffer exchanged to 20 mM Tris, pH 7.5, 600 mM NaCl for crystallization needs. For MAXI-scale, all reagents were scaled linearly.

### Protein purification using a 3XFLAG-tag

Purification using a 3XFLAG-tag has been carried out using ANTI-FLAG M2 magnetic beads from Sigma-Aldrich. All magnetic separations were done using the MagneSphere technology Magnetic Separation Stand from Promega. Per 1.2 ml translation (MIDI-scale), 170 μl of gel suspension (85 μl of packed gel) was used. The needed volume of gel suspension was first equilibrated in TBS buffer and then combined with the DTT-free translation mixture. The binding was carried out for 1 hour at room temperature with shaking on the ThermoMixer C (~1000 rpm). Using the magnetic stand, the supernatant was removed and discarded, and the beads were then washed 3 times with TBS buffer (0.5 ml each time). To elute the bound protein, TBS buffer supplemented with 150 μg/ml 3XFLAG peptide (purchased from Sigma) was applied to the beads, and the mixture was left shaking for 20 min. Then, the supernatant fraction was collected, and the elution procedure was repeated one more time. Two elution fractions were combined and concentrated to ~ 100 μl using 10kDa or 30 kDa amicons from EMD Millipore. For MAXI- and MEGA-scale reactions, reagents were scaled linearly but final concentrated volume remained around 100 μl.

To remove the 3XFLAG tag, a light chain enterokinase from New England Biolabs, NEB, was employed. Due to inhibition of this enzyme at high concentrations of salts and its requirement for the Ca^2+^ ions, ~100 μl of purified and concentrated protein was further buffer-exchanged using the Bio-Rad spin mini columns, filled with Bio-gel P-6DG (pre-soaked in 20 mM Tris, pH 7.5, 50 mM NaCl and 2 mM CaCl_2_). Because the enterokinase cleavage efficiency is time- and template concentration-dependent, small-scale cleavage titration was carried out first to determine the right conditions: 10 μl of concentrated protein was mixed with 2 μl of 1.6 U/μl enterokinase. Aliquots (2 μl) were taken over time up to 3 hours and the cleavage efficiency was analyzed by running the samples on the SDS-PAGE. Once the appropriate time was determined, the rest of the protein was cleaved in a similar fashion. For single particle analysis, the removal of enterokinase was not required and thus the sample was buffer exchanged back to TBS using Bio-gel P-6DG (pre-soaked in TBS), split in aliquots, frozen in liquid nitrogen and then transferred to −80°C for a long-term storage. In case where prolonged protein incubation was later required at ambient temperatures either for crystallization or assembly trials, the enterokinase was removed. For enterokinase removal, the reaction was stopped with 1/10^th^ volume of 5M NaCl and mixed with 15 μl if 100% pre-equilibrated agarose-trypsin slurry from Sigma. The mixture was further incubated at room temperature for 30 minutes at 300 rpm on the ThermoMixer C. Upon completion, the mixture was directly applied on spin mini column, containing Bio-Gel P6-DG (presoaked in TBS buffer). The flow-through was collected, quantified, frozen in liquid nitrogen and transferred to −80°C for a long-term storage.

### PAGE electrophoresis

Protein expressions were evaluated on 10% Mini-PROTEAN TGX Precast Protein gels from Bio-Rad. For SDS-PAGE, 3 μl of a protein solution was mixed with 4 μl of 500 mM DTT and 7 μl of the 2X SDS Sample Buffer. Then the sample was denatured at 90°C for 5 minutes, snap-cooled on ice and loaded on the gel. The general running conditions for SDS were 150V in the 1X Tris-Glycine-SDS Running Buffer from Bio-Rad. The gels were further stained with Bio-Safe Coomassie Stain from Bio-Rad. Molecular ladders used were Broad Range Protein Molecular Weight Markers from Promega and Kaleidoscope^™^ Prestained Standards (#161-0324 and #161-0375) from Bio-Rad. Native PAGE was performed using the same 10% Mini-PROTEAN TGX Precast Protein gels where 12 μl of protein solution was mixed with 12 ul of 2X Native Sample Buffer from Bio-Rad prior loading (no heat denaturation employed).

To analyze the proteins comprising FluoroTect GreenLys, the denaturation protocol for SDS-PAGE was changed to 70°C for 2 minutes, and a protein loading per well was increased to 6 μl. After the run was complete, the fluorescent images of gels were collected immediately by FluorChem Q from Alpha Innotech or a laser-based scanner Typhoon FLA 9500 from GE Healthcare, set for Alexa488 excitation and emission.

### 2D protein crystallization using a Ni^2+−^chelating lipid monolayer

Purified CCMK protein with 6xHis tag on C-terminal (15 μl, 0.3 mg/ml) was adjusted with imidazole to 10 mM and glycerol to 10% and loaded in the concave well of PELCO Immunostaining Pad from Ted Pella. Lipid mixture was prepared by combining 4 parts of 10 mg/ml of DOPC (in chloroform) with 1 part of 2.5 mg/ml of 18:1 DGS-NTA(Ni) (in chloroform), achieving the final weight ratio of 16:1. Both lipids were purchased from Avanti Polar Lipids. To create a monolayer, 2 μl of lipid mixture was deposited on the air-water interface of the protein solution. The PELCO pad was then enclosed inside the sealed Petri dish, containing wet filter papers to create a humid environment and prevent evaporation. The 2D crystallization was carried out at room temperature for 13 hours.

### TEM imaging and data analysis

Negative staining of proteins (PDX1.2, SEO1, GS proteins) was carried out in the following manner: 1) an ultrathin carbon film on lacey carbon (01824, Ted Pella) was glow discharged at 10 mA for 30 seconds using PELCO easiGlow from Ted Pella, 2) 3 μl of protein solution was deposited on the carbon side and let to absorb for 1 minute, 3) the excess of solution was blotted away, and the grid was floated on a 20 μl drop of the NanoW stain from Nanoprobes for 1 min, 4) then the excess of the stain was wicked away, and the grid was let to air-dry. For 2D protein crystals (CCMK protein), an ultrathin carbon film on lacey carbon grid was not glow-discharged. It was directly dropped carbon side down on top of lipid layer at the air-water interface of protein solution. The grid was then lifted and transferred to a drop of Nano-W stain, incubated for 1 minute, blotted and air-dried. The EM micrographs were acquired on 300kV FEI Environmental TEM, 300 kV FEI Scanning TEM and 300kV JEOL JEM-3000SFF. For CCMK crystals, the Focus suite has been used for image processing, FFT determination, lattice unbending, and mapping [50]. For cryo-EM experiments, C-flat holey grids (CF213, EMS) were used instead. Prior to sample deposition, they were first glow-discharged at the same conditions as above and then transferred to the plunge-freezer Leica EM GP, brought to 80% humidity at 25 °C. Then 3 μl of PDX 1.2 protein sample (50 μg/ml, in TBS buffer) was pipetted on the carbon side and blotted for 3 s before plunge-freezing in the liquid ethane. The frozen samples were then transferred to liquid nitrogen and then to 300kV JEOL JEM-3000SFF, pre-cooled with liquid helium to 4K. The images were collected at low-dose conditions at defocus values of 2-4 μm using DE20 camera from Direct Electron. Collected images were further processed using CisTEM software [51], installed on local supercomputer.

### X-ray crystallization trials

Cell-free produced GS protein (tagged with 3XLAG, 10 mg/ml, 100 μl) was crystallized in hanging drop format by the addition of 0.2 M sodium chloride, 0.1 M BIS-Tris pH 5.5 and 25% w/v PEG 3350. The TTP Labtech Mosquito nanodispenser was used to set up 210 nl drops of protein and reservoir solution mixed in a 1:1, 1:2, and 2:1 ratio. Drops equilibrated over 100 ul of reservoir solution in a 96-well tray. Crystals appeared over several days. Crystals were soaked for ten seconds in a 2:1 solution of mother liquor and glycerol.

## Acknowledgements

This work was supported by DOE-BER Mesoscale to Molecules Bioimaging Project FWP# 66382. A portion of the research was performed using the Environmental Molecular Sciences Laboratory (EMSL), a national scientific user facility sponsored by the Department of Energy’s Office of Biological and Environmental Research and located at PNNL. X-ray crystallization trials were carried out at X-ray Crystallography Core facility at UCLA, supported by DOE Grant DE-FC02-02ER63421. Diffraction data was obtained at the Northeastern Collaborative Access Team beamlines, which are funded by the National Institute of General Medical Sciences from the National Institutes of Health (P41 GM103403). The Eiger 16M detector on 24-ID-E beam line is funded by a NIH-ORIP HEI grant (S10OD021527). This research used resources of the Advanced Photon Source, a U.S. Department of Energy (DOE) Office of Science User Facility operated for the DOE Office of Science by Argonne National Laboratory under Contract No. DE-AC02-06CH11357.

## Author contributions

JEE managed all aspects of this work. JEE and IVN devised experiments for the study. IVN performed genetics, molecular biology, cell-free expression and electron microscopy experiments. NS performed cryo-EM screening, data collection and analysis of PDX sample. TM performed all work with CCMK sample. RS assisted IVN with material handling. MJC and DC performed x-ray crystallization screen and diffraction data collection for GS sample. YL, HH, and MK provided clones of PDX and SEO genes and assisted with analysis of the resulting proteins. IVN and JEE wrote initial manuscript draft but all authors contributed to writing the manuscript and approval of final version.

## Supplemental Material

**Figure S1.**
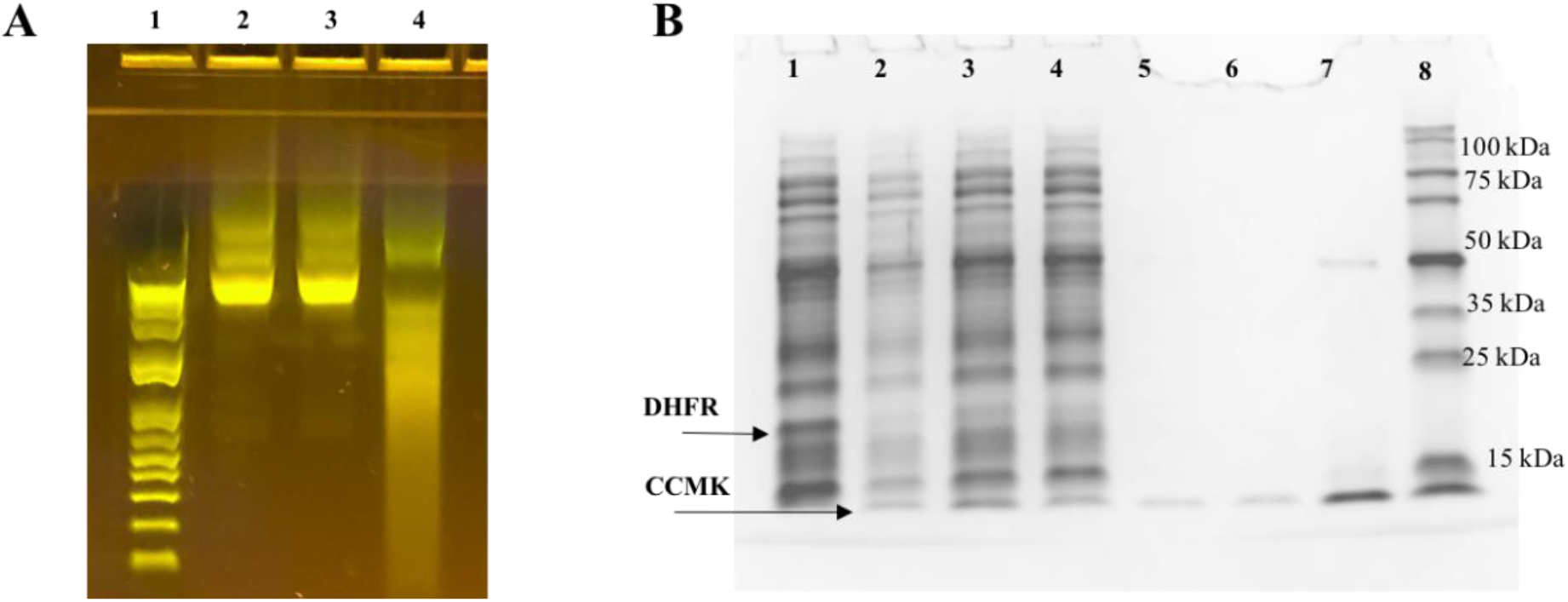
A. The detection of various mRNAs on agarose. Lane 1, MW DNA markers; Lanes 2-4, various mRNA products. B. SDS-PAGE gel, showing the expression and purification of CCMK protein using 6xHis-tag and wash buffer supplemented with 10 mM imidazole. Lane 1, control DHFR expression; Lane 2, crude mixture; Lane 3, soluble fraction; Lane 4, flow-through; Lane 5, elution fraction 1; Lane 6, elution fraction 2; Lane 7, elution fractions 1 and 2 combined and concentrated; Lane 8, MW markers.

**Figure S2.**
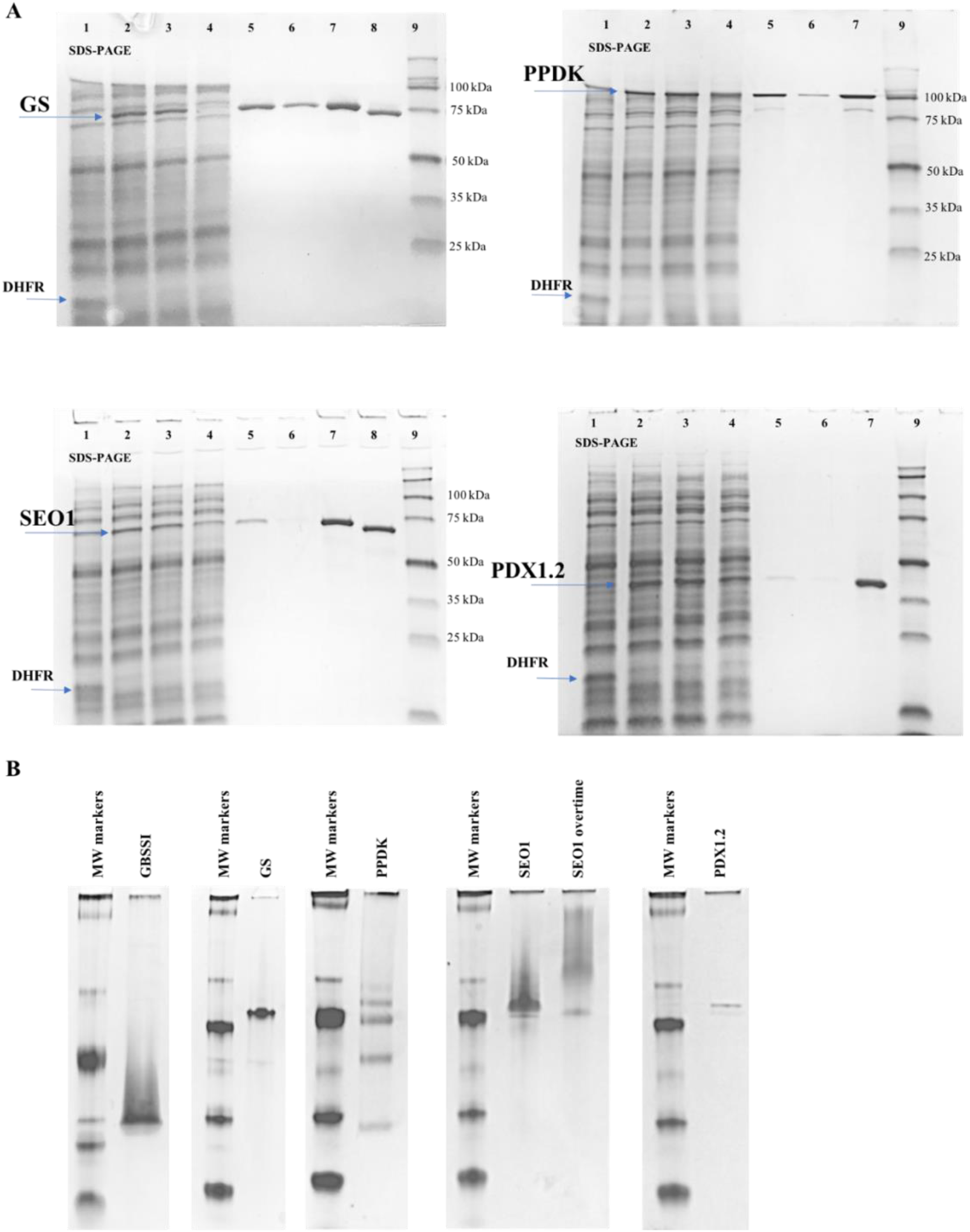
Protein purification and analysis by PAGE. A. SDS-PAGE gel of expression and purification of various proteins using 3XFLAG-tag. Lane 1, control DHFR expression; Lane 2, crude mixture; Lane 3, soluble fraction; Lane 4, flow-through; Lane 5, elution fraction 1; Lane 6, elution fraction 2; Lane 7, elution fractions 1 and 2 combined and concentrated; Lane 8, 3XFLAG tag removed; Lane 9, MW markers. B. Native PAGE analysis

**Figure S3.**
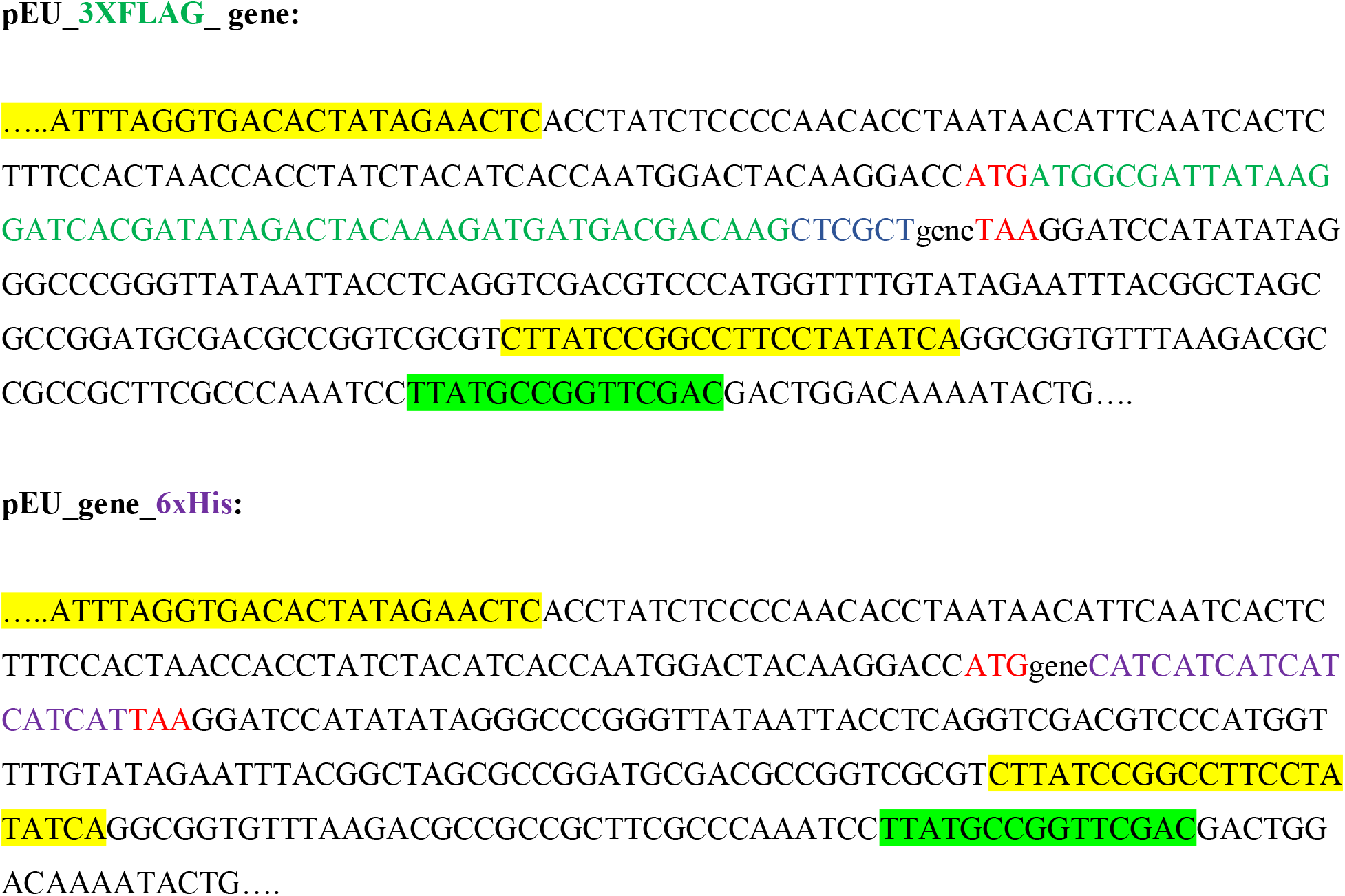
The sequence design of pEU constructs used in this study.

